# Comparing lipid remodeling in mouse adipose and liver tissue with quantitative Raman microscopy

**DOI:** 10.1101/2022.03.07.483297

**Authors:** Alexandra Paul, Belén Chanclón, Cecilia Brännmark, Pernilla Wittung-Stafshede, Charlotta S. Olofsson, Ingrid Wernstedt Asterholm, Sapun H. Parekh

**Affiliations:** Department of Biology and Biological Engineering, Division of Chemical Biology, Chalmers University of Technology, Kemivägen 10, 412 96 Gothenburg, Sweden; Department of Biomedical Engineering, University of Texas at Austin, 107 W Dean Keeton Street, Austin, TX 78712, USA; Department of Physiology (Metabolic Physiology), Institute of Neuroscience and Physiology, Sahlgrenska Academy at the University of Gothenburg, Box 432, 405 30 Gothenburg, Sweden; Department of Molecular Spectroscopy, Max Planck Institute for Polymer Research, Ackermannweg 10, 55 128 Mainz, Germany

**Author notes:** Corresponding author. S. H. Parekh, Department of Biomedical Engineering, University of Texas at Austin, 107 W Dean Keeton Street, Austin, TX 78712, USA.

**Keywords:** adipose tissue, diet, mouse, Raman spectroscopy, triglyceride

## Abstract

Brown adipose tissue (BAT) consists of highly metabolically active adipocytes that catabolize nutrients to produce heat. Playing an active role in triacylglycerol (TAG) clearance, research has shown that dietary fatty acids can modulate the TAG chemistry deposition in BAT after weeks-long dietary intervention, similar to what has been shown in white adipose tissue (WAT). Our objective was to compare the influence of sustained, non-chronic dietary intervention (a one-week interval) on WAT and BAT lipid metabolism and deposition *in situ*. We use quantitative, label-free chemical microscopy to show that one week of HFD intervention results in dramatically larger lipid droplet (LD) growth in BAT (and liver) compared to LD growth in inguinal WAT (IWAT). Moreover, BAT showed lipid remodeling as increased unsaturated TAGs in LDs, resembling the dietary lipid composition, while WAT (and liver) did not show lipid remodeling on this time scale. Concurrently, expression of genes involved in lipid metabolism, particularly desaturases, was reduced in BAT and liver from HFD-fed mice after one week. Our data show that BAT lipid chemistry remodels exceptionally fast to dietary lipid intervention compared WAT, which further points towards a role in TAG clearance.

## 1. INTRODUCTION

Lipids are efficient energy storage molecules and have evolved in their use as energy repositories to bridge times of energy scarcity. In the developed world, most humans struggle not with food scarcity, but rather, they have essentially unlimited access to nutrients, which has led to a brimming obesity epidemic that is escalating worldwide (1). A constant positive energy balance (largely resulting from excess nutrient ingestion) leads to increased body weight associated with an increased risk for the development of metabolic diseases such as type 2 diabetes and cardiovascular disease, as well as certain cancers (2) and the sever of upper respiratory infections (3). Excess energy can be stored as lipids *via* two distinct pathways: (i) *de novo* lipogenesis from glucose or amino acids or (ii) direct esterification of non-esterified fatty acids (NEFAs). Both pathways will result in formation of neutral lipids, primarily triacylglycerols (TAGs), which are stored in lipid droplets (LDs) located predominantly in adipose tissue (4). White adipose tissue (WAT) TAGs derive mostly from direct esterification of NEFAs that are formed either *de novo* in the liver or taken up from the diet (5). WAT is also responsible for supplying NEFAs to the other organs to meet the body’s energy demand (6). Importantly, the balance between uptake and release of NEFAs needs to be tightly controlled, to match the body’s energy needs. Excess fatty acids that are stored in the adipose tissue can lead to ectopic lipid deposition associated with chronic inflammation and insulin resistance (7, 8). Conversely, increased energy expenditure and/or effective storage of excess nutrients in the form of neutral lipids protects against metabolic disturbances.

Searching for possibilities to increase energy expenditure to reduce obesity and its co-morbidities, brown adipose tissue (BAT) has emerged as a potential therapeutic target (9–11). BAT is the key site for non-shivering thermogenesis, a process crucial for the ability of small rodents and hibernating animals to keep a constant body core temperature. This heat production is achieved in the mitochondria, which contain uncoupling protein-1 (UCP1) that bypasses the respiratory chain to generate heat instead of ATP from fatty acid oxidation. NEFAs from TAGs are the main fuel for heat production in rodent BAT (12). Studies of BAT in high fat diet (HFD)-fed rodents showed that BAT mass and lipid content increase during weight gain in mice, indicating that the TAG accumulation in BAT exceeds the catabolic processes (13, 14). Interestingly, increased lipid deposition can be observed within the first day of HFD feeding, which shows the acute reaction of BAT to increased lipid intake (15). It was also shown that the TAG chemistry in BAT changes dramatically during cold acclimation and arousal from hibernation, highlighting the plasticity in lipid chemistry in this tissue (16). While the lipid storage role of BAT is still being clarified, the lipid storage function of WAT has been well established as accumulating TAGs in direct response to dietary lipids. That is, the TAG composition in WAT ultimately correlates with the composition of ingested dietary lipids (17, 18). However, the adaptation of lipid depots in BAT and WAT (and liver) – particularly in terms of lipid chemistry – to a relatively short lipid-rich dietary intervention has not been compared.

We designed a study to compare the impact the TAG clearing ability of BAT and WAT, in terms of LD morphology and TAG chemistry. We measure the molecular composition of TAGs and gene expression profiles in BAT, inguinal WAT (IWAT), gonadal WAT (GWAT, supplementary information), and liver in mice in response to one week of HFD intervention and compare to chow-fed littermates. For the characterization of TAGs, we used *in situ*, label-free lipid chemical imaging from quantitative coherent Raman microscopy, which can be used to accurately determine lipid biochemistry (chain length and number of double bonds) with sub-micrometer resolution and has been validated against the gold-standard of mass spectrometry (19, 20). Combined the with additional characterization of lipid metabolism gene expression, we found that HFD diet challenge for as little as one-week results in differences in lipid uptake / *de novo* lipogenesis and mRNA expression between WAT and BAT (as well as liver) tissues.

## 2. MATERIALS AND METHODS

### 2.1. Mice and diets

All animal experiments were performed after approval from the local Ethics Committee for Animal Care at the University of Gothenburg, Sweden, and followed appropriate guidelines. Male C57BL/6J mice were obtained from Charles River Laboratories at four weeks old and allowed to acclimate in the animal facility for one week under standard housing conditions with *ad libitum* access to food and water. For the one-week intervention, littermates were randomly selected to be either kept on a standard laboratory chow diet (CD, Special Diets Services, N=5) or on a high fat diet (HFD, D12492, Research Diets Inc., N=5). Mice were fasted for 4h prior to tissue collection. Blood glucose levels were measured using a glucometer (Contour from Bayer; Baser, Switzerland). Dietary fatty acid composition, mice body weight, and blood glucose levels at the end point can be found in **Supplementary Figures S1** and **S2**. BAT, IWAT, GWAT, and liver were snap-frozen in liquid nitrogen after extraction.

### 2.2. Tissue preparation for microscopy

For combined CARS and SHG microscopy whole tissue (BAT, IWAT, GWAT, or liver) was fixed in cold 4% para-formaldehyde (PFA) containing 150 mM sucrose at 4°C while the tissues thawed and then mounted between a glass cover slide and cover slip (#1) submerged in PBS. For BCARS analysis frozen adipose tissue (IWAT, GWAT, or BAT) was sliced at −39°C in a MTC cryotome (Slee Medical GmbH) using low profile diamond blades (C.L. Sturkey Inc.) into 20 μm thick slices, while liver was cut at −14°C and slices were 10 μm thick. The tissue slices were directly transferred to chilled class cover slips, immediately covered with 4% PFA containing 150 mM sucrose in the cryotome chamber, sandwiched with a second cover slip, and placed in a refrigerator at 4°C. Afterwards, all cover slide/slip sandwiches were sealed with nail polish before further analysis, and all imaging was done at room temperature.

### 2.3. Coherent anti-Stokes Raman scattering and second harmonic generation microscopy

Lipids and collagen were visualized in intact, thawed tissue pieces using a SP 5 II TCS CARS microscope (Leica) equipped with a pico-second pulsed laser source at 817 and 1064 nm (Pico Emerald, APE). Tight focusing conditions were achieved with a water immersion objective (L 25x, NA=0.95, Leica) and 2845 cm^-1^ CARS signals for lipids were collected above the sample via a CARS 2000 filter set in front of a photomultiplier tube. SHG of collagen was collected in epi direction *via* a 390/40 bandpass filter in front of a photomultiplier tube.

### 2.4. Lipid droplet and cell volume determination

The CARS images of all tissues showing the LDs in a single optical section were thresholded using the default method in ImageJ and a watershed algorithm was employed to separate LDs. A region of interest was selected manually to only include areas where LDs were perfectly separated by the water shedding. Then, the ImageJ “particle analysis” function was used to calculate LD area. LD volume was retrieved assuming a perfect sphere to calculate the volume from the area measured in the center of the LD (as manually chosen from a z stack). Between three and 10 images were analyzed per mouse and tissue. From that volume, the average LD diameter was calculated. Cell volumes for BAT are manually estimated from intrinsic fluorescence of the cells at high contrast settings (n=50 per condition). The cell size was not analyzed for WAT due to the low cytosolic volume and neither for liver due to the unclear cell borders.

### 2.5. Broadband CARS (BCARS) microscopy

The home-built setup with custom software written in LabVIEW (National Instruments) has been described in detail before (21). Briefly, a super continuum Stokes beam was generated *via* a photonic crystal fiber after a dual-output laser source (Leukos-CARS, Leukos), which also provided a narrow band pump/probe beam (λ = 1064 nm). The Stokes beam was passed through a short pass filter and Glan-Thompson polarizer producing a bandwidth from 1100-1600 nm with a power density of 100 μW nm^-1^. It was then combined with the pump/probe beam in the focal plane of an inverted microscope (Eclipse Ti-U, Nikon) *via* an air immersion objective (100x, NA 0.85, Zeiss). Raster scanning was achieved with a XYZ piezo stage (Nano-PDQ 375 HS, Mad City Labs). The BCARS signals were collected with a 10x air immersion objective (NA 0.25, Zeiss) above the sample, where the laser beams were removed with a notch (NF03-532/1064E-25, Semrock) and short-pass filter (FES1000) before the signal was analyzed by a spectrometer (Shamrock 303i, Andor) with a CCD camera (Newport DU902P-BR-DD, Andor). The collected spectra ranged from 900-3500 cm^-1^ with a spectral pitch of ~ 4 cm^-1^. Images were collected with 500 nm step size and 15-20 ms pixel dwell for adipose tissues and 250 nm steps with 100 ms pixel dwell for liver. For all samples a reference spectrum was collected in glass. Two spectral images were collected per mouse and tissue.

### 2.6. BCARS data analysis

The imaginary component of the third-order Raman susceptibility from the raw BCARS spectra was extracted with a modified Kramers-Kronig algorithm (22). This was followed by error-phase correcting employing an iterative noise-maintaining approach that is model-free *via* the use of a second-order Savitzky-Golay filter (101 spectral points, 404 cm^-1^) (23). The resulting Raman-like spectra were fitted pixel by pixel to generate maps of TAG chain length and saturation as described elsewhere (19). After the initial fit, an additional threshold of 12-24 carbons #C=C > −0.1 was applied to ensure a physiological range. On average, less than 1% of pixel were discarded in this step, typically at droplet borders where lipid intensity in the spectra was too low to be fitted accurately. The 3×3 images (101×101 pixel) were then stitched and false color coding was applied to display TAG chain length and #C=C.

### 2.7. Total RNA extraction and quantitative real-time PCR

Total RNA from BAT, IWAT, GWAT, and liver was extracted using the TissueLyser LT (Qiagen) and the RNeasy^®^ Lipid Tissue Mini Kit (Qiagen) including an on-column DNase digest (RNase-Free DNase Set, Qiagen) according to manufacturer’s instructions. RNA concentration and purity were determined with a NanoDrop Spectrometer. A total of 500 ng RNA per sample were used for cDNA preparation (High-Capacity cDNA Reverse Transcription Kit, Applied Biosystems) according to the manufacturer’s instructions. Quantitative real-time PCR was performed using Fast SYBR^®^ Green Master Mix (Applied Biosystems) and the QuantStudio™ 7 Flex System. Primer sequences are listed in **Supplementary Table 1**. 5 ng cDNA was amplified per sample and all samples were amplified in duplicates under nuclease-free conditions. Conditions were 95°C for 20 s (1 cycle), 95°C for 1 s, 60°C for 20 s, 72°C for 15 s (40 cycles), followed by a melting curve (0.5°C / s) from 60°C to 95°C to confirm one PCR-product. Gene-expression was calculated as relative expression to β actin (*Actb*) via 2^ΔCT^.

### 2.8. Statistical analyses

LD volume and 2^ΔCT^ values were averaged per mouse. To compare average TAG chain lengths and number of double bonds between the two dietary conditions, all chain lengths/number of double bonds from two large-area (150×150 μm^2^) scans per mouse were averaged. Statistical testing was then performed between the two groups of mice. Then a two-tailed, unpaired Mann-Whitney test was performed comparing each HFD intervention with its respective CD control using GraphPad Prism 6. P<0.05 was considered significant. P values for the gene expression analysis are listed in **Supplementary Table 2**. Values are typically represented by their mean with standard deviation (SD).

## 3. RESULTS

### 3.1. Experimental Outline

To induce obesity, we fed male C57CL/6J mice (24) either CD or HFD with 7 or 60% caloric intake from lipids, respectively (**Supplementary Figure S1A**). The CD supplied 4 kcal% mixed sugars while the HFD contained 6.8 kcal% sucrose. On average, the diets supplied NEFAs with 17.4 (CD) /17.5 (HFD) carbons and 1.3 (CD) / 1.0 (HFD) double bonds per lipid chain (**Supplementary Figure S1B, C**). Although there was a slight difference in saturation in the two diets, both contained more average #C=C than the average TAG chains we found in all investigated tissues (detailed below). The HFD intervention lasted one week and a significant increase in body weight was observed compared to the CD fed mice with similar circulating glucose levels, as expected for mice with the genetic background (25) (**Supplementary Figure S2**).

### 3.2. One week of HFD induced changes in LD morphology and increases TAG unsaturation in BAT

In all tissue samples (BAT, WAT, and liver), LDs were easily identified by their strong signal in label-free CARS vibrational imaging at an energy of 2845 cm^-1^. This imaging highlights CH2 groups, which are abundant in lipid acyl chains; we and others have used this methodology extensively in lipid imaging (19, 21, 26, 27). The lipid droplet and cell morphology were investigated in intact, fixed tissue with minimal sample preparation (no embedding or staining). In BAT, we observed that the volume of individual LDs increased almost 40-fold after one week of HFD as compared to CD controls (**Figure 1A, B**; LD volume distribution: **Supplementary Figure S3A**). LDs from HFD BAT were on average 11.1 ± 0.8 μm (mean ± SD, CD 3.2 ± 0.2 μm) in diameter after one week (**Supplementary Figure S3A**) and adipocytes in BAT from HFD-fed mice appeared to lose their multilocular LD morphology and shift towards a morphology with one large central LD per cell (**Figure 1A**, top left images), resembling that of white adipocytes. Interestingly, the initial increase of LD volume did not lead to an increase in cell size (based on 2-photon autofluorescence) compared to the multilocular CD cells (**Figure 1C**). Using broadband coherent Raman microscopy (BCARS), we analyzed the biochemistry (chain length and number of double bonds) of TAGs in LDs. Our algorithm allows us to map TAG chain length and number of double bonds on a pixel-by-pixel basis (19). We report the average chain length and unsaturation per fatty acid chain in TAGs over all TAG molecules within a focal volume (300 nm x 300 nm x 5000 nm). We have previously shown this approach to show absolute accuracy of 0.7 carbons and 0.2 double bonds when compared to GC-MS for more than 10 different oils, with the advantages of mapping TAG chemistry *in situ* and virtually no sample preparation (19).

**Figure 1:**
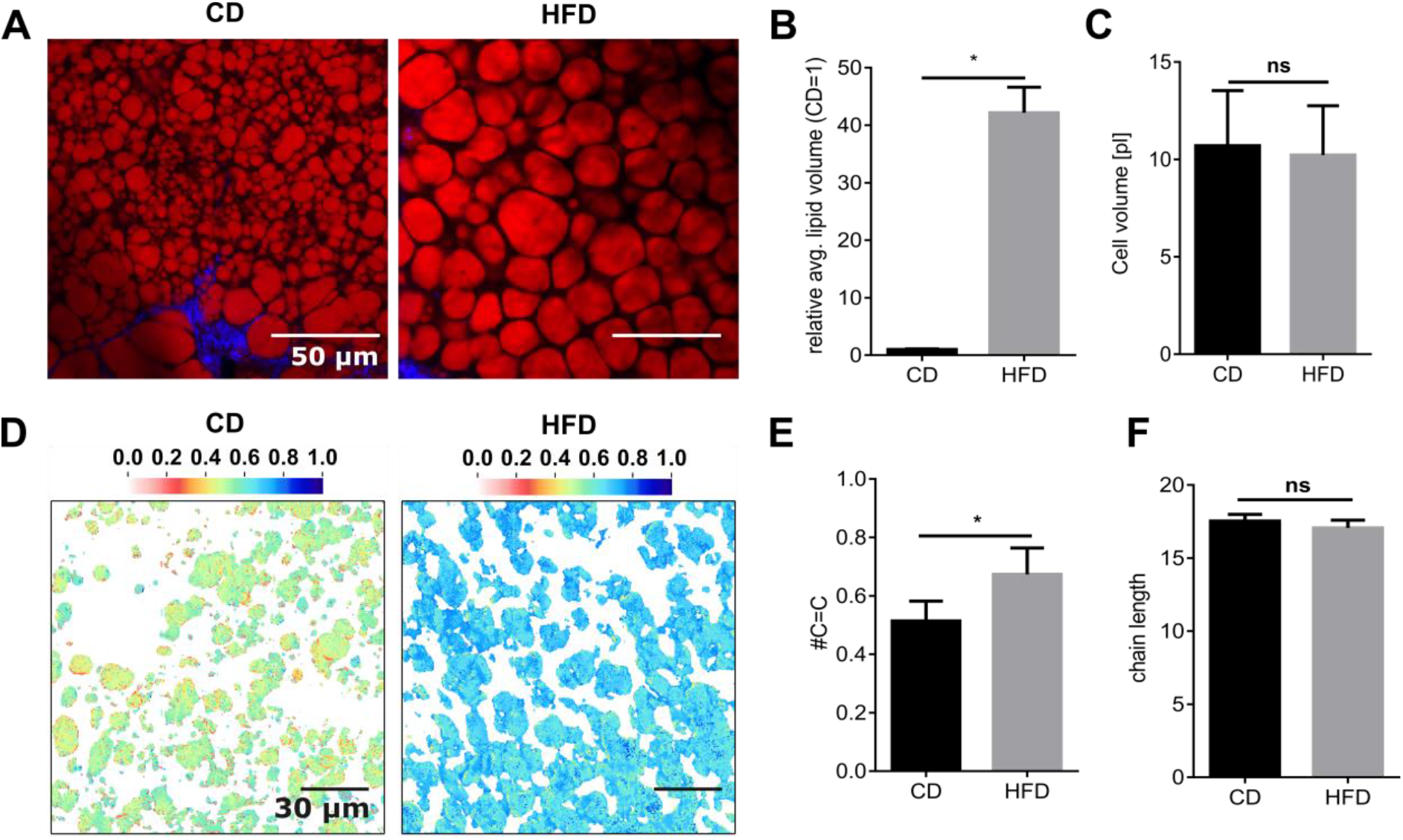
HFD-induced changes in BAT. **A,** The size of lipid droplets under high fat diet compared to chow diet (red – lipids, CARS 2845 cm^-1^; blue – SHG, collagen, scale bar: 50 μm) increases visibly and quantitatively as seen by the **B,** average volume per droplet in comparison to the chow diet (normalized to 1). Statistical analysis is done between absolute values: p=0.0159. **C,** At the same time, the cells in BAT do not increase in size. **D,** Images of the average number of double bonds per TAG chain in BAT LDs for CD (left) or after one-week HFD (right). **E,** Average number of double bonds of the TAGs inside the LDs increases significantly after one week of HFD feeding in BAT; p=0.0079. **F,** Average chain length of the TAGs inside the LDs is not affected. Values are displayed as mean ± SD. * P < 0.05 and ** P < 0.01 by multiple two-tailed, unpaired Mann-Whitney tests.

From BCARS chemical imaging, we found the number of double bonds – reflecting unsaturation – showed a significant increase after one week of HFD as compared to CD (**Figure 1E**) and no changes in TAG chain length in the LDs after HFD (**Figure 1F**). Chemical imaging also showed that the intracellular TAG unsaturation uniformly increased one week of HFD feeding (**Figure 1D**).

### 3.3. One-week HFD feeding increased LD volume in IWAT and liver while TAG chemistry was unchanged

As in BAT, we mapped the lipid distribution in IWAT using single-frequency CARS microscopy. We found LD hypertrophy after one week of HFD in IWAT (**Figure 2A, B**; LD volume distribution: **Supplementary Figure S3B**), while the lipid chemistry was not affected (**Figure 2C-F**). The lipid unsaturation distribution between the one-week CD and HFD diets are nearly indistinguishable, in contrast what we found for BAT (**Figure 1D**).

**Figure 2.**
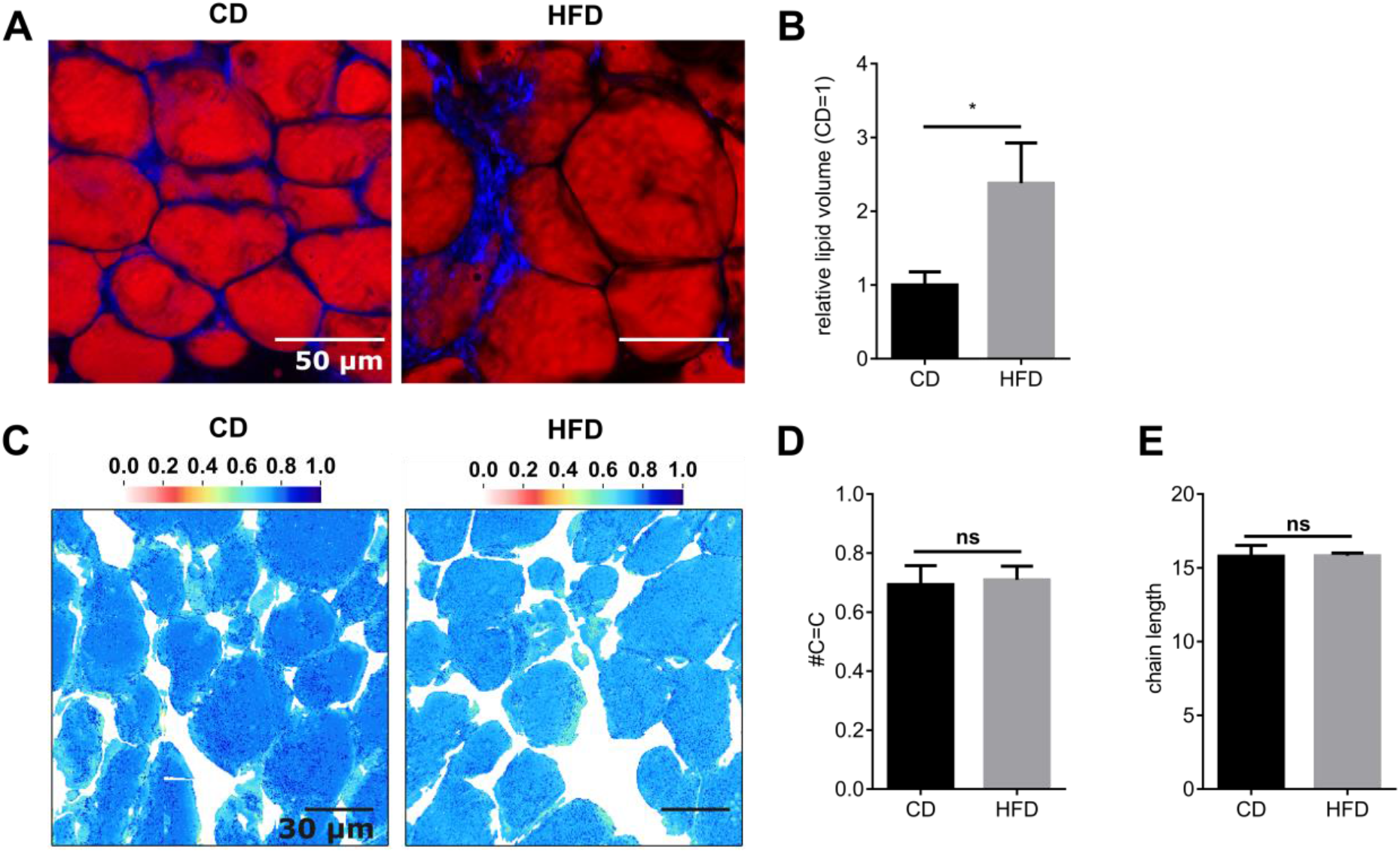
HFD-induced changes in IWAT. **A,** IWAT shows a less drastic increase in LD size after one week of HFD compared to CD (red – lipids, CARS 2845 cm^-1^; blue – SHG, collagen) as seen by **B,** the average lipid volume per droplet (normalized to CD=1). Statistical analysis is done between absolute values: p=0.0317. **C,** Images of the average number of double bonds per TAG chain in IWAT LDs for CD (left) or after one-week HFD (right). **D,** Average number of double bonds per TAG chain is statistically identical under CD or one week of HFD feeding. **E,** Average lipid chain length in TAGs is statistically identical after one week of HFD feeding in IWAT. Values are displayed as mean ± SD. * P < 0.05 and ** P < 0.01 by multiple two-tailed, unpaired Mann-Whitney tests.

Notably, while all investigated adipose tissues (BAT and WAT) seemed to have stored similar chain lengths (15-17 carbons), the amount of double bonds differed vastly between tissues. Adipocytes in BAT from CD-fed mice contained the lowest abundance of double-bond-containing TAGs (0.53±0.05, mean ± SD). In response to one week of HFD, TAGs in BAT showed increased double bonds, reaching a level of unsaturation found in IWAT from chow-fed (or HFD) mice (#C=C bonds in HFD (BAT) = 0.69±0.07; #C=C bonds in CD(IWAT) = 0.68±0.04). This demonstrates a differential remodeling of the BAT vs. WAT lipid droplets to the same one-week dietary intervention. We note that gonadal WAT (GWAT) showed no difference between the two diets in LD size or TAG chemistry (**Supplementary Figure S4**). The lower number of double bonds in BAT in mice on the CD highlights the different metabolic state of the two adipose depots.

We next investigated how lipid content and chemistry changed with HFD intervention in the liver since the liver is an important organ in lipid metabolism. Interestingly, we found a drastic (7-fold) increase of LD volume in liver after one week of HFD (**Figure 3A, B**; LD volume distribution: **Supplementary Figure S3D)**. As in all BAT tissues, LDs in liver displayed lower # of double bonds compared to the CD in WAT; however, the increase in unsaturation after one week on the HFD did not reach statistical significance (**Figure 3C, D**). The average chain length was not affected by diet (**Figure 3E**). In comparison to all other investigated tissues, liver LDs showed the highest inter-LD heterogeneity (**Figure 3C**), particularly for mice on the CD.

**Figure 3.**
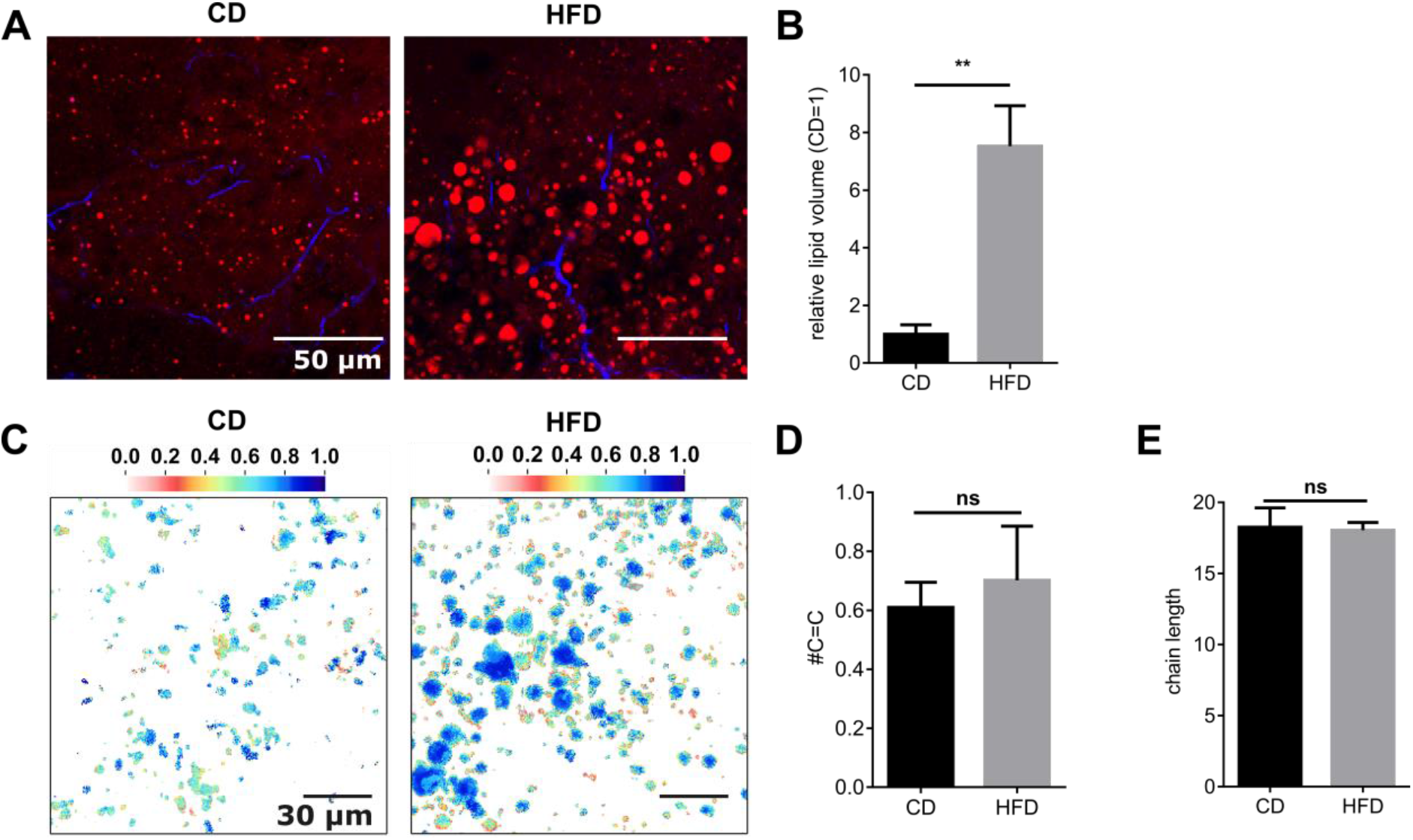
HFD induced changes in liver. **A,** Liver shows an increase in LD size after one week of HFD compared to CD (red – lipids, CARS 2845 cm^-1^; blue – SHG, collagen) as seen by **B,** the average lipid volume per droplet (normalized to CD=1). Statistical analysis is done between absolute values: p=0.0079. **C,** Images of the average number of double bonds per TAG chain in liver LDs for CD (left) or after one-week HFD (right). **D,** Average number of double bonds per lipid TAG chain is statistically identical under CD or one week of HFD feeding. **E,** Average lipid chain length in TAGs is statistically identical after one week of HFD feeding in liver. Values are displayed as mean ± SD. * P < 0.05 and ** P < 0.01 by multiple two-tailed, unpaired Mann-Whitney tests.

### 3.4. Genetic changes in BAT, WAT, and liver show reduced *de novo* lipogenesis

To further probe the mechanism of increased HFD-induced unsaturation in BAT, we compared mRNA expression of numerous genes involved in *de novo* lipogenesis and lipid modification, particularly fatty acid desaturases (**Figure 4A**). We found that the expression of genes involved in *de novo* lipogenesis in BAT, *e.g*., desaturases *Scd1* and *Scd2*, elongase (*Elovl6*) and fatty acid synthase (*Fasn*) was significantly reduced after one of HFD, as compared to CD. Moreover, *Agpat2* (1-Acylglycerol-3-Phosphate O-Acyltransferase 2), an enzyme catalyzing the second step of *de novo* phospholipid biosynthesis, was also reduced after one week of HFD intervention compared to the CD controls. The regulator of sterol biosynthesis, *Srebf1c* (Sterol Regulatory Element Binding Transcription Factor 1), was not affected by the HFD. The expression of genes involved in TAG synthesis, *Gpam* (Glycerol-3-Phosphate Acyltransferase, Mitochondrial), *Dgat1/2* (Diacylglycerol O-Acyltransferase 1/2), and *Gpd1* (Glycerol-3-Phosphate Dehydrogenase), was not affected. The expression of *Ucp1* (uncoupling protein 1), related to non-shivering thermogenesis, mitochondrial biosynthesis (*Ppargc1a* - Peroxisome proliferator-activated receptor gamma coactivator 1-alpha), *Lpl* (lipoprotein lipase), and gluconeogenesis/glyceroneogenesis (*Pck1* - Phosphoenolpyruvate Carboxykinase 1) were not significantly different in the HFD mice than in the CD mice. *Cpt1a/b* (Carnitine Palmitoyltransferase 1B), involved in beta-oxidation, was also unchanged. Taken together, the gene expression analysis suggests that *de novo* lipogenesis was largely suppressed by HFD intervention even after one week, which indicates that the changes in lipid unsaturation in BAT LDs was due to increased dietary NEFA uptake. This is supported by the isotope tracing of lipids in BAT 2h post feeding (**Supplementary Figure S6**).

**Figure 4.**
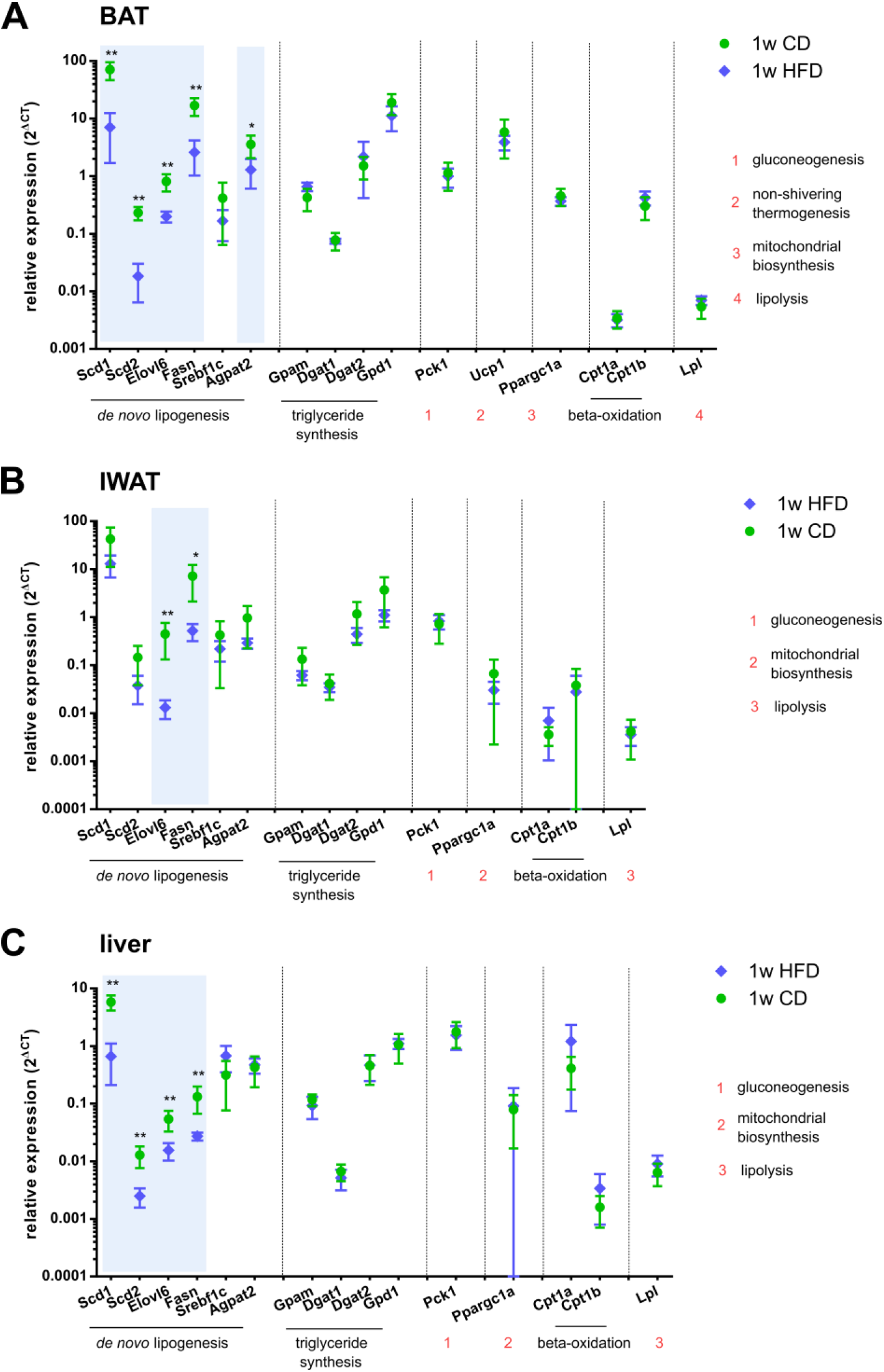
Diet induced gene expression changes. Relative gene expression (2^ΔCT^) compared to β actin in **A,** BAT, **B,** IWAT, and **C,** liver (green circles – CD, blue diamond – HFD). Significant downregulation in comparison to CD are marked with stars and shaded in blue. All p values can be found in **Supplementary Table 2**. N=5. Values are displayed as mean ± SD. * P < 0.05 and ** P < 0.01 by multiple two-tailed, unpaired Mann-Whitney tests.

In IWAT, mRNA levels of all genes involved in *de novo* lipogenesis, *Elovl6* and *Fasn*, were reduced after the one-week HFD intervention (**Figure 4B**) while all other genes remained unaffected. In GWAT most genes in the *de novo* lipogenesis pathway were also significantly reduced after one week (**Supplementary Figure S5**).

Our findings of increased fatty acid content and concurrent downregulation of genes linked to *de novo* lipogenesis in IWAT and BAT after HFD suggest increased uptake of NEFAs that are than stored as TAGs in in those tissues. NEFAs can come from either the diet or liver metabolism, where NEFAs and TAGs are regularly synthesized, signifying the importance of the liver gene expression. Genes involved in the production of longer and more unsaturated chains, *Scd1/2, Elovl6*, and *Fasn*, were downregulated after one-week of HFD analogous to BAT. No other genes were differentially expressed after one week (**Figure 4C**). Once again, it appears that the surplus of available NEFAs from the diet correlates with lower *de novo* synthesis.

## 4. DISCUSSION

In this study, we explored the changes in TAG chemistry in lipid-containing tissues induced by HFD intervention for one week. As a complementary approach to studies that rely on radioactive labeling or extraction of all lipids, we investigated both cell and LD morphology and TAG chemistry in intact tissue sections as well as expression of genes involved in various lipid metabolic pathways in the examined tissues. While our lipid quantification method cannot resolve the identity or abundance of individual lipids such as in mass spectrometry, it can accurately map both TAG chain length and #C=C with sub-μm precision unraveling inter- and intra-cellular differences with almost no sample preparation.

### 4.1. HFD induced unilocular LDs in BAT and increases in LD size in all investigated tissues

HFD challenges the whole body with a constant oversupply of lipids that, under isocaloric conditions, would principally be used as energy or stored in WAT depots. However, if the amount of supplied lipids exceeds that which can be stored in WAT, ectopic lipid deposition in non-adipose tissues, like the liver, will occur. These ectopic depositions can arise from long-term lipid-rich diets (7), but also as a mechanism to protect the rest of the body during short term overfeeding (28, 29). Our study focused on a relatively short, sustained intervention (one week), and we used average LD volume to judge the amount of stored lipids in adipose and non-adipose tissues. Our data showed that liver LD volume grew significantly, by 7.5-fold over CD control, after one week (**Figure 3B**). LD volume in BAT showed an even higher increase, 42-fold, after one week of HFD. After the HFD dietary intervention, cells in BAT displayed unilocular LDs instead of the multilocular LDs found under CD (**Figure 1A** and **Figure 5**). This again implies that BAT is highly responsive to an acute lipid surplus, as has been shown previously (15). The increase in large LDs in BAT after HFD intervention was linked to a ‘whitening’, coupled to loss of BAT functionality in other studies (30). However, we did not observe any reduction in the mRNA expression of *Ucp1* after HFD intervention, suggesting that BAT still functions in non-shivering thermogenesis. Contrary to BAT and liver, the relative change in IWAT LD volume was smaller with no change in GWAT LDs (**Figure 2B, Supplementary Figure S4**), highlighting the different response of consumptive versus storage tissues. This suggests that BAT may be the primary protector against lipid overload during short duration high-fat intake whereas WAT may be more important for longer sustained dietary intake of high fat.

**Figure 5.**
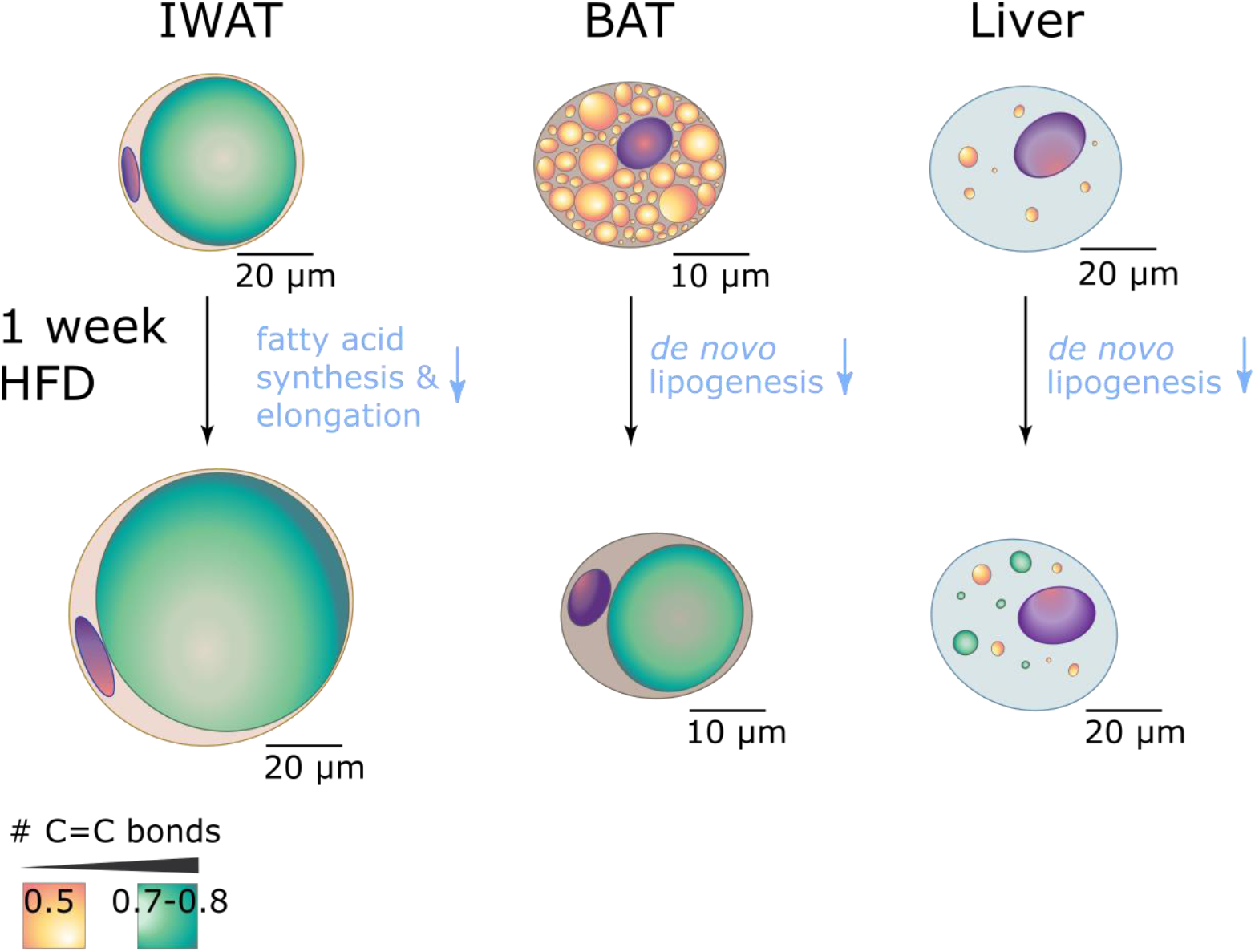
Schematic of cellular changes in IWAT, BAT, and liver. The color of the LDs in the cells represents the average number of C=C bonds (orange 0.5 C=C, green 0.7-0.8 C=C). Increases in average LD/cell volume are to scale. Pathways where gene expression is reduced are marked in blue. The average LD volume is increased in all tissues after one week of HFD feeding and adipocytes in BAT display more unilocular LDs. The increase in TAGs correlates with reduced activity in the *de novo* lipogenesis pathway. However, only BAT shows a uniform significant increase in LD TAG unsaturation.

### 4.2. More unsaturated TAGs are found in LDs during HFD feeding in BAT after one week

While HFD increased LD size in all studied tissues, the fatty acids stored as TAGs in BAT showed a marked increase in #C=C bonds. In comparison to the #C=C in extracted tissues (0.43-0.89 as seen using the minimum to maximum values over all tissues from our data), the # of C=Cs supplied with the diet were 1.0 (HFD). We found that the TAG unsaturation increased with HFD in BAT compared to the CD. After the same intervention of HFD feeding, we found that the TAG chemistry was not significantly affected in IWAT or liver. There are three possible reasons why BAT showed more prominent changes in LD chemistry: 1) reduced *de novo* lipogenesis and increased uptake of dietary NEFAs, 2) a starting chemical state of TAGs in BAT that was more saturated, or 3) preferential uptake of unsaturated fatty acids in BAT. Our data with isotope labelled triolein, with 1 C=C bond per chain, showed that BAT took up more than 4-fold triolein compared to WAT and liver, highlighting a fast clearance of dietary lipids by BAT. This is in line with results reported by Bartelt *et al.* describing the rapid clearance of dietary fatty acids by BAT specifically (31). Combined with the mRNA expression data showing decreased *de novo* lipogenesis, this supports reason 1. Next, our BCARS data show that TAGs in BAT LD indeed start with fewer C=C bonds, on average, compared to WAT or liver. With a 60% caloric intake on the HFD consisting of unsaturated lipids with 1.0 C=C bond per chain and fast clearance of lipids by BAT, there is “more room” to increase unsaturation in BAT in support of reason 2. Finally, it is also possible that BAT shows preferential uptake of unsaturated dietary fatty acids compared to saturated fatty acids. Previous work has shown that monounsaturated and n-6 polyunsaturated fatty acids were preferably taken up from the diet in the postprandial period for IWAT (32) in support of reason 3. Therefore, a combination of reasons 1, 2, and 3 could lead to the changes in TAG chemistry in BAT LDs with one-week HFD.

Turning to WAT, we found no changes in LD unsaturation, suggesting a slower lipid turnover compared to BAT. This is supported by the comparatively low isotope triolein uptake (**Supplementary Figure S6**). Our findings on WAT are consistent with that of Brunengraber *et al.* that showed incorporation of dietary fatty acids into GWAT in male C57BL/6J mice such that the relative abundance of unsaturated chains (C18:1, C18:2) in the TAGs increased after 34 days compared with 5 days (33). In liver, TAGs in LDs from the one-week CD and HFD showed more heterogeneity both in terms of chain length and #C=C bonds compared to WAT or BAT. Such inter-LD heterogeneity in hepatocytes for TAG, cholesterol content, and associated perilipins has been described previously (34).

### 4.3. BAT and liver showed early reduction of genes involved in de novo lipogenesis or lipid chemical modification

In addition to imaging changes in TAG chemistry, we compared mRNA expression of genes involved in modifying TAG chemistry, as well as other genes of interest for TAG storage or utilization. There was a minor difference in sugars supplied by the diets (4 % kcal (CD), 6.8 % kcal (HFD)), which could potentially drive *de novo* lipogenesis during HFD. However, we found no differences in *Pck1* (encoding for PEPCK-C the rate-limiting enzyme in gluconeogenesis (35)) in all adipose tissues or liver after HFD compared to CD feeding, suggesting an equal contribution of gluconeogenesis/glyceroneogenesis with both diets. In WAT, uptake of circulating NEFAs from diet or uptake from *de novo* lipogenesis from the liver is the dominating source of lipids (5). Liver is the main contributor to *de novo* lipogenesis (36, 37). In all investigated tissues, the fatty acid synthesis pathway (measured by *Fasn* expression) was significantly reduced after one week of HFD intervention. This is in line with previous findings that HFD significantly downregulates the contribution of *de novo* lipogenesis upon one week of high-fat feeding in rats (38).

As genetic markers of TAG chemical remodeling, we looked at fatty acid elongase (*Elovl6*) and desaturase enzymes (*Scd1* and *Scd2*), which can modify fatty acid chain length and saturation, respectively. We found *Elov16* in liver and BAT to be significantly downregulated after one week of HFD while the TAG chain lengths were unchanged. HFD-induced downregulation of both *Scd1* and *Scd2* were observed in comparison to CD after one week in liver and BAT. If the TAGs inside the LDs would be from *de novo* lipogenesis, the reduced activity of these key genes would lead to a decrease in desaturation and chain length. However, this trend is not supported by our data on LD chemistry.

## 5. CONCLUSIONS

Our study highlights the different metabolic timescales of BAT, WAT, and liver tissues in response to one-week HFD challenge. We measured LD size and chemistry *in situ* in BAT, WAT, and liver isolated from HFD-fed or CD-fed mice using quantitative, label-free chemical microscopy. Connecting these measurements to genetic changes in the same tissues, we found that dietary intervention has distinct effects on BAT and liver compared to white adipose tissues. BAT, often considered an oxidative tissue, showed an increase in TAG #C=C and larger increase in LD size in response to HFD intervention after one week than WAT, the classical lipid storage tissue. Gene expression analysis supported the notion that dietary fatty acids contributed to lipid remodeling, with HFD inducing a reduction in expression of genes related to *de novo* lipogenesis in BAT and liver. Our work suggests that acute uptake of fatty acids is buffered in in BAT (and to some extent liver), which not only changes LD size but also composition.

## ACKNOWLEDGEMENTS

We thank Frederik F. Fleissner for support with the BCARS setup. The research leading to these results has received funding from the European Union’s Seventh Framework Program (FP7/2007-2013) under grant agreement n°607842, Kungl. Vetenskaps-och Vitterhets-Samhället, Wilhem och Martina Lundgrens Stiftelse, Marie Curie Foundation n°CIG322284, Swedish Research Council (2013-7107, 2017-00792, 2019-00682 and 2020-01463), Novo Nordisk Foundation (NNF19OC0056601), Swedish Diabetes Foundation, Welch Foundation (F-2008-20190330), and the Human Frontiers in Science Program (RGP0045/2018).

