## Supplementary Information for "Comparing lipid remodeling in mouse adipose and liver tissue with quantitative Raman microscopy"

### Content

*Supplementary Tables*

**Table S1. Primer sequences (5’→3’).**

**Table S2. Summary of p-values**.

*Supplementary Figures*

**Figure S1. Dietary compositions**.

**Figure S2. Mice body weight and blood glucose**.

**Figure S3. Lipid droplet volume distributions.**

**Figure S4. HFD-induced changes in GWAT.**

**Figure S5. Diet induced gene expression changes in GWAT.**

**Figure S6. Postprandial lipid uptake in CD-fed mice.**

*Supplementary Methods*

**Method S1. Lipid uptake.**

*Supplementary Abbreviations*

BAT – brown adipose tissue, CD – chow diet, GWAT – gonadal white adipose tissue, HFD – high fat diet, IWAT – inguinal white adipose tissue,

*Supplementary Tables*

**Table S1. Primer sequences (5'→3').**

| Gene | forward | reverse |
|---|---|---|
| *Actb* | GACCCAGATCATGTTTGAGA | GAGCATAGCCCTCGTAGAT |
| *Scd1* | TTCTTGCGATACACTCTGGTGC | CGGGATTGAATGTTCTTGTCGT |
| *Scd2* | GCATTTGGGAGCCTTGTACG | AGCCGTGCCTTGTATGTTCTG |
| *Elovl6* | GAAAAGCAGTTCAACGAGAACG | AGATGCCGACCACCAAAGATA |
| *Fasn* | GGAGGTGGTGATAGCCGGTAT | TGGGTAATCCATAGAGCCCAG |
| *Srebf1c* | ATCGGCGCGGAAGCTGTCGGGGTAGCGTC | ACTGTCTTGGTTGTTGATGAGCTGGAGCAT |
| *Agpat2* | CGTGTATGGCCTTCGCTTTG | TCCATGAGACCCATCATGTCC |
| *Gpam* | ACAGTTGGCACAATAGACGTTT | CCTTCCATTTCAGTGTTGCAGA |
| *Dgat1* | TCCGTCCAGGGTGGTAGTG | TGAACAAAGAATCTTGCAGACGA |
| *Dgat2* | CTGGCTGATAGCTGCTCTCTACTTC | TGTGATCTCCTGCCACCTTTC |
| *Gpd1* | GTGAGACGACCATCGGCTG | TTGGGTGTCTGCATCAGGT |
| *Pck1* | CTGCATAACGGTCTGGACTTC | CAGCAACTGCCCGTACTCC |
| *Ucp1* | GTGAAGGTCAGAATGCAAGC | AGGGCCCCCTTCATGAGGTC |
| *Ppargc1a* | TATGGAGTGACATAGAGTGTGCT | CCACTTCAATCCACCCAGAAAG |
| *Cpt1a* | TGGTGGGAGGAATACATC | CAGAAGACGAATAGGTTTGAG |
| *Cpt1b* | GCACACCAGGCAGTAGCTTT | CAGGAGTTGATTCCAGACAGGTA |
| *Lpl* | GGTTGCGCGTAGAGAGGATG | CTCACGCTCTGACATGCCTTC |

**Table S2. Summary of p-values** for two-sided, unpaired Mann-Whitney test comparing $2^{\Delta CT}$ (β actin reference) values of HFD against CD. Blue indicates significant down regulation compared to the CD.

| | BAT | IWAT | GWAT | liver |
|---|---|---|---|---|
| | **1 week** | **1 week** | **1 week** | **1 week** |
| | HFD vs CD | HFD vs CD | HFD vs CD | HFD vs CD |
| **de novo lipogenesis** | | | | |
| *Scd1* | 0.0079 | 0.1508 | 0.1508 | 0.0079 |
| *Scd2* | 0.0079 | 0.0556 | 0.0079 | 0.0079 |
| *Elovl6* | 0.0079 | 0.0079 | 0.0079 | 0.0079 |
| *Fasn* | 0.0079 | 0.0159 | 0.0079 | 0.0079 |
| *Srebf1c* | 0.2222 | 0.5317 | 0.0079 | 0.0952 |
| *Agpat2* | 0.0317 | 0.1508 | 0.0159 | 0.8016 |
| **triacylglyceride synthesis** | | | | |
| *Gpam* | 0.0566 | 0.6667 | 0.5317 | 0.1508 |
| *Dgat1* | 0.3095 | 0.4127 | 0.6667 | 0.2222 |
| *Dgat2* | 0.9444 | 0.4127 | 0.2222 | 0.9444 |
| *Gpd1* | 0.1508 | 0.1508 | 0.2222 | 0.5317 |
| **gluconeogenesis** | | | | |
| *Pck1* | 0.8016 | 0.6667 | 0.0556 | 0.5317 |
| **non-shivering thermogenesis** | | | | |
| *Ucp1* | 0.5317 | | | |
| **mitochondrial biosynthesis** | | | | |
| *Ppargc1a* | 0.4127 | | | |
| **beta-oxidation** | | | | |
| *Cpt1a* | 0.9444 | 0.4127 | 0.4127 | 0.0952 |
| *Cpt1b* | 0.3076 | 0.8016 | 0.0159 | 0.2222 |
| **lipolysis** | | | | |
| *Lpl* | 0.3095 | 0.9444 | 0.5317 | 0.3095 |

<u>*Supplementary Figures*</u>

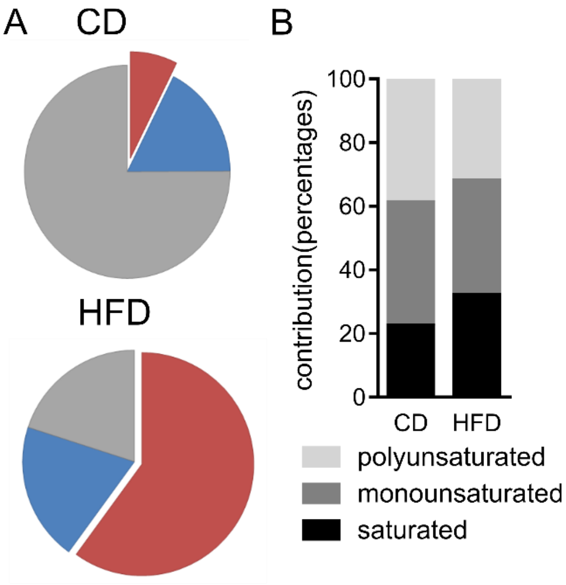


C

| | Chow Diet | High Fat Diet |
|---|---|---|
| **Saturated Fatty Acids** | *22.5* | *32.0* |
| C10:0 Capric | | 0.04 |
| C12:0 Lauric | 0.9 | 0.1 |
| C14:0 Myristic | 6.2 | 1.1 |
| C15:0 Pentadecylic | | 0.1 |
| C16:0 Palmitic | 13.7 | 19.6 |
| C17:0 Margaric | | 0.4 |
| C18:0 Stearic | 1.8 | 10.6 |
| C20:0 Arachridic | | 0.2 |
| **Monounsaturated Fatty Acids** | *38.8* | *36.0* |
| C14:1 Myristoleic | 0.9 | |
| C16:1 Palmitoleic | 4.0 | 1.3 |
| C18:1 Oleic | 33.9 | 34.0 |
| C20:1 Paullinic | | 0.6 |
| **Polyunsaturated Fatty Acids** | *38.8* | *32.0* |
| C18:2(ω6) Linoleic | 30.4 | 28.7 |
| C 20:2 Eicosadienoic | | 0.8 |
| C18:3(ω3) Linolenic | 2.6 | 2.0 |
| C20:3 Dihomo-γ-linolenic acid | | 0.1 |
| C20:4(ω6) Arachidonic | 5.7 | 0.3 |
| C22:5(ω3) Clupanodonic | | 0.1 |
| **Average chain length** | **17.43** | **17.56** |
| **Average #C=C** | **1.30** | **1.03** |

**Figure S1. Dietary compositions. A,** Contributions to 100% daily caloric intake from carbohydrates (grey), protein (blue), and lipids (red) for chow (CD, top) or high fat diet (HFD, bottom. **B and C,** Fatty acid composition (in %) in the 7 or 60 % lipid for CD or HFD, respectively.

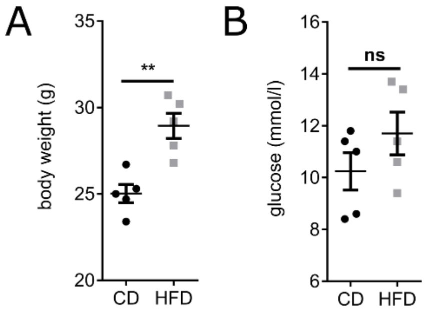


**Figure S2. Mice body weight and circulating glucose levels. A,** Weight gain and **B,** circulating glucose levels after 1-week (N=5, p=0.0079) HFD intervention. Mice were fasted for 4h prior to glucose measurements and tissue collection. Values are displayed a mean (SD). Two-tailed, unpaired Mann-Whitney test.

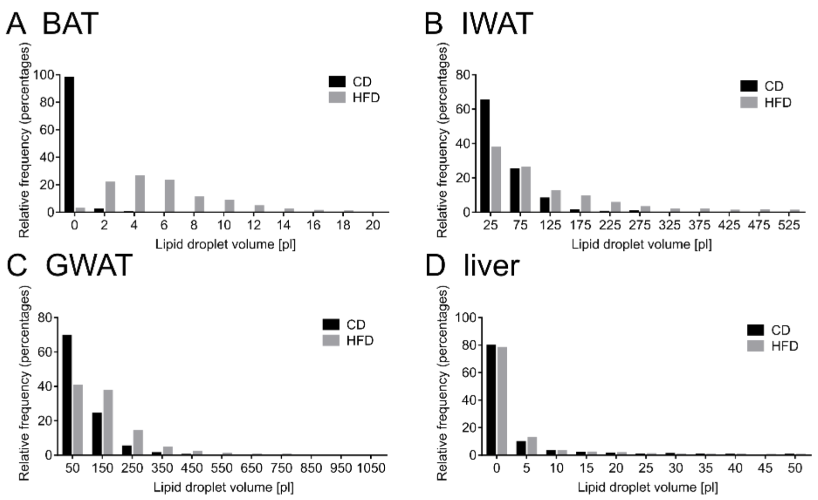


**Figure S3. Lipid droplet volume distributions.** Binned histograms of all lipid droplets in **A)** BAT (CD n=3236, HFD n=392), **B)** IWAT (CD n=337, HFD n=567), **C)** GWAT (CD n=527, HFD n=679), and **D)** liver (CD n=1869, HFD n=1416). N= 5 (1week). For analysis procedure see Methods 2.4. Due to different lipid droplet volumes in the different tissues, the bin sizes were adapted to 2 pl BAT/liver, 50 pl IWAT, and 100 pl GWAT. Each tissue was normalized by the total lipid droplet number to calculate percentages per bin.

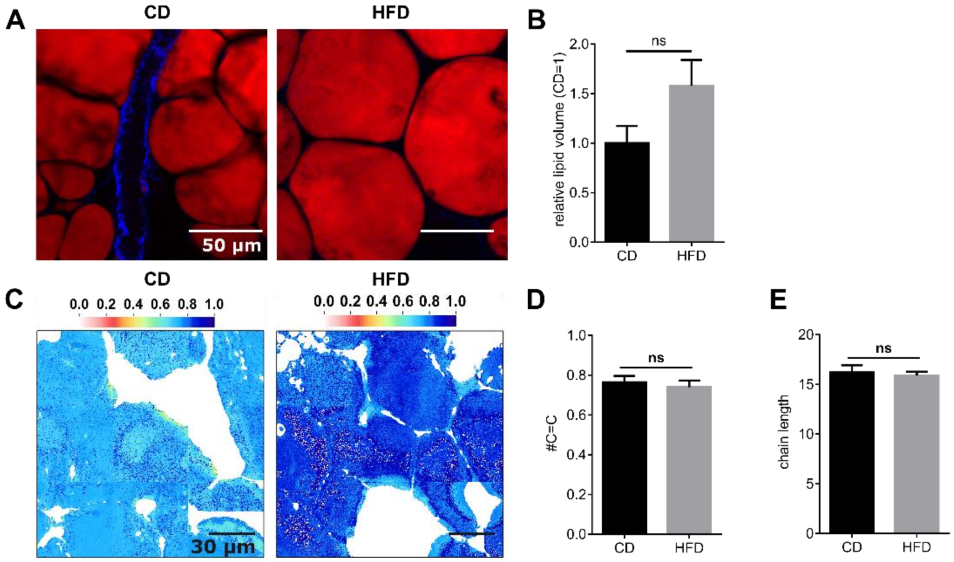


**Figure S4. HFD-induced changes in GWAT. A,** GWAT shows no significant increase in LD size after one week of HFD compared to CD (red – lipids, CARS 2845 cm-1; blue – SHG, collagen) as seen by **B,** the average lipid volume per droplet (normalized to CD=1). Statistical analysis is done between absolute values. **C,** Images of the average number of double bonds per TAG chain in GWAT LDs for CD (left) or after one-week HFD (right), **). D,** Average number of double bonds per lipid TAG chain is statistically identical under CD or one week of HFD feeding. **E,** Average lipid chain length in TAGs is statistically identical after one week of HFD feeding in GWAT. Values are displayed as mean ± SD. * $P < 0.05$ and ** $P < 0.01$ by multiple two-tailed, unpaired Mann-Whitney tests.

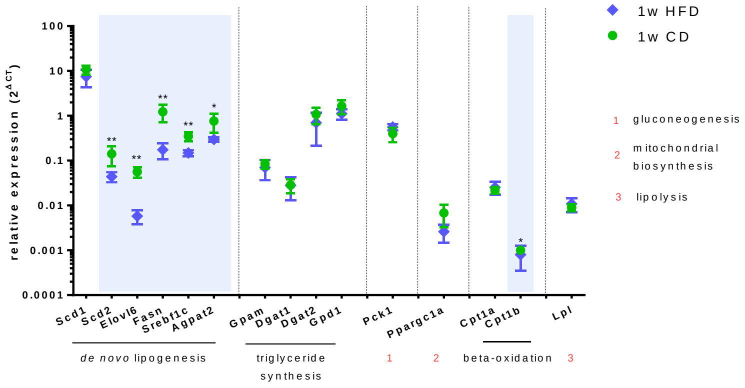


**Figure S5. Diet induced gene expression changes.** Relative gene expression ($2^{\Delta CT}$) compared to β actin in GWAT (green circles – CD, blue diamond – HFD). Significant downregulation in comparison to CD are marked with stars and shaded in blue. All p values can be found in **Supplementary Table 2**. N=5. Values are displayed as mean ± SD. * P < 0.05 and ** P < 0.01 by multiple two-tailed, unpaired Mann-Whitney tests.

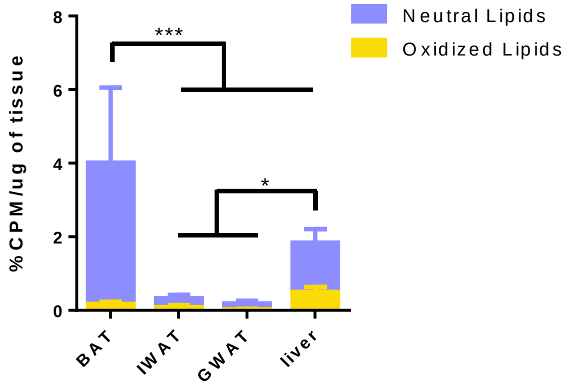


**Figure S6. Lipid uptake in CD-fed mice. A,** After 2h BAT shows a significant increased uptake of radioactively labeled neutral lipids in CD-fed mice (N=8, p<0.0001). **B,** When comparing the uptake of all lipids BAT takes up the largest amount (N=8, p≤0.0004), while liver takes up more than both WATs (N=8, p≤0.0175). Values are displayed a mean (SD). 2way ANOVA with Tukey's multiple comparisons test. For analysis procedure see Supplementary Methods S1.

*Supplementary Methods*

**Method S1. Lipid uptake.**

Animals and diet design

All animal experiments were performed after approval from the local Ethics Committee for Animal Care at the University of Gothenburg, Sweden, and followed appropriate guidelines. Male C57BL/6J mice were obtained from Charles River Laboratories (Germany) and housed under standard conditions (12 hours light/dark) with ad libitum access to water and CD diet (Special Diet Services, UK, full diet composition in Supplementary Figure S1A). Mice (N=8) were kept on CD for 12 weeks before the lipid uptake test was performed.

Lipid uptake

A radioactive labelled tracer was used to assess the lipid uptake in the different tissues (BAT, IWAT, GWAT, liver). The tracer (Triolein, [9,10-3 H(N)], Perkin Elmer, Boston, MA, USA) was incorporated at 2μCi/mouse into a 20% intralipid emulsion (300 ul/mouse; Sigma-Aldrich, St Louis, MO, USA). After 4h fasting, the mice were given an oral load by gavage. After 2h, tissues were collected in 2:1 chloroform:methanol, homogenized using a tissue lyser, and stored at 4C over night. A 1M $CaCl_2$ solution was added to all samples, centrifuged at 3000rpm (4C, 20min), separating the aqueous phase (oxidized fraction) from the organic phase (accumulated fraction). The phases were separated into scintillation vials with 5ml Ultima Gold™ scintillation cocktail (Perkin Elmer, USA). The amount of incorporated 3H was measured in a beta counter (Perkin Elmer, USA), corrected to the mass of the tissue, and presented as percentage of the total ingested volume per mouse.
